# Superoxide Dismutase 1 translocates to the nucleus in spinal cord mixed cells from newborn rats

**DOI:** 10.1101/2022.03.08.483426

**Authors:** Nathália V. Santos, Camila M. L. Machado, Giselle Cerchiaro

## Abstract

Superoxide dismutase 1 (SOD1) catalyzes the superoxide conversion to oxygen and hydrogen peroxide in general. SOD1 can translocate to the cell nucleus in response to oxidative stress in yeast and human fibroblasts. Here, we report the translocation of sod1 to the cell nucleus in primary co-cultures of neurons and astrocytes derived from newborn rats explants even in the absence of oxidative stress stimuli. The successful tissue explants from rats allowed simplistic and clean modeling of Amyotrophic lateral sclerosis (ALS)-simile tissue microenvironment. This mixed-cells-population was responsive to H_2_O_2_ (100 μM) oxidative stress with a progressive dying process up to 6 h after stimuli. However, no differences in SOD1 nucleus translocation were observed in rat amyotrophic lateral sclerosis (ALS)-simile tissue microenvironment up to 6 hours under oxidative stress. These results altogether point to a robust cellular collaboration between neurons and astrocytes so that the tissue can equilibrate this SOD1 translocation during the oxidative stress response and in the early animal development.

## 1. Introduction

Reactive oxygen species (ROS) are the products of healthy metabolism and xenobiotic exposure, and depending on their concentration, ROS can be beneficial or harmful to cells and tissues (Circu and Aw, 2010; Halliwell, 1992; Miller et al., 2019; Poprac et al., 2017; Valko et al., 2007). ROS include, predominantly, the superoxide anion (O_2_^-^), hydrogen peroxide (H_2_O_2_), hydroxyl radical (OH^•^), and nitrogen-oxygen derived radicals (Fridovich, 1978)(Augusto et al., 2002; Medinas et al., 2007). At low physiological levels, ROS function as messengers in intracellular signaling and regulation, whereas excess ROS induce oxidative modification of cellular macromolecules and induces cell death (Sies and Jones, 2020)(Finkel, 2011)(11). Evolutionarily, several antioxidant strategies were selected for cells to deal with ROS toxicity. Among the agents considered as antioxidants, enzymes catalytically remove reactive radicals and species, such as the enzymes glutathione reductase, glutathione peroxidase, catalases and superoxide dismutase (SOD) (Al-Kayiem and Ibrahim, 2015; Lei et al., 2015).

The superoxide dismutases class enzymes, SODs, have the role of scavengers of superoxide, present in prokaryotes, and eukaryotes, they have the function to protect the cells from endogenously generated superoxide anions during aerobiosis (Fridovich, 1995). SODs are metalloenzymes that are broadly classified based on the metal cofactor used. Based on their binding of the metal cofactor, SODs are classified into four types: manganese co-factored (MnSOD), iron co-factored (FeSOD), copper-zinc co-factored (Cu-ZnSOD or SOD1) and nickel co-factored (NiSOD) (Miller, 2012)(Culotta et al., 2006). These different types of SODs are localized in separate cellular compartments, and the MnSOD is mainly present in mitochondria and peroxisomes. FeSOD has been mostly detected in chloroplasts and peroxisomes, and Cu-Zn-SOD or SOD1 has been detected in the cytosol, chloroplasts, and peroxisomes (Sheng et al., 2014)(Zelko et al., 2002). The compartmentalization of different forms of SOD throughout organisms allows them to counteract various stresses locally. SOD out-competes damaging reactions of superoxide, thus protecting the cell from superoxide toxicity. Besides serving as a critical antioxidant for detoxification of the radicals that are usually produced within cells, SOD has been shown to be significant in disease development (Che et al., 2016; Fukai and Ushio-Fukai, 2011). Cells within tissue collaborate to sustain tissue homeostasis during different injuries as oxidative stress. Some authors attribute the neurodegeneration observed in Amyotrophic Lateral Sclerosis (ALS) to imbalances in oxidative stress (Shibata et al., 2001).

Here we took advantage of two *in vitro* models to report data linking SOD1 nuclear translocation during oxidative stress in astrocytes and neurons cells. We show cell death and cellular translocation of SOD1 during oxidative stress caused by H_2_O_2_ in a mixed astrocytes and motor neuron primary cell culture (ALS-simile tissue microenvironment) derived from spinal cord rat explants. However, no-differences on SOD1 nuclear translocation were observed in the ALS-simile tissue microenvironment, suggesting a potential robust equilibrium between astrocytes and motor-neuron during an oxidative stress-injury.

## 2. Materials and methods

### 2.1 Chemicals

Unless otherwise stated, chemicals were obtained from Sigma-Aldrich and were of analytical grade: solutions were prepared using Milli-Q water (Millipore).

### 2.2 Cell Cultures

The cell media were made with DNase- and RNase-free water and filtered through 0.22-μm filter membranes (Millex GV, Millipore) before use. The cell cultures were manipulated using sterile, disposable non-pyrogenic plastic ware and were maintained at 37 ºC in an atmosphere of 5% CO_2_ in air at a relative humidity of 80%.

### 2.3 Extraction and Isolation of spinal cord cells

The primary spinal cord cell culture procedure described was based on a previously described method (Gingras et al., 2007). Post-natal day 0-2 rats were decapitated following the local animal ethical committee rules and approving and were carried out under the U.K. Animals (Scientific Procedures) Act, 1986 and associated guidelines. The skin and muscles overlying the spinal cord were dissected. The spinal cord between the cervical and spinal cord was removed and maintained in a sterile cold PBS solution. The spinal cord was washed with sterile, cold PBS containing 100 U/mL penicillin and 10.0 μg/mL streptomycin (Sigma) and cut into small fragments. The tissue was incubated in trypsin solution for 15 min a 37°C and triturated with a Pasteur pipette. The cells were pelleted at 4000 rpm for 5 minutes, the supernatant was removed, and the pellet resuspended in 5mL of cold PBS. Purification of spinal cord cells was performed using OptiprepTM (Sigma), which is a 10.4% solution of Optpret in PBS. To separate spinal cord cells, 5mL of the cell suspension in PBS was layered slowly onto the 10.4 % solution of Optiprep and centrifuged at 1900 rpm for 20 min at room temperature. The spinal cell fraction was carefully collected as a first band at the interface of the PBS and the 10.4% solution of Optiprep. That fraction was pooled into a centrifuge tube containing PBS and centrifuged at 2000 rpm for 5 min.

### 2.4 Culture of spinal cord cells

The cells were resuspended in Neurobasal A medium containing 100U/mL penicillin and 100 μg/mL streptomycin (Sigma) and supplemented with 25 μg/mL gentamycin (Sigma), L-glutamine (Invitrogen), GlutaMAX™ (Invitrogen), B-27® Supplement (Invitrogen) and 10% of fetal bovine serum (Gibco). The cells were plated into culture plates coated with 0.01% cell culture-tested polylysine solution (Sigma, molecular weight: 150,000–300,000 Da), and maintained at 37 °C in of 5 % CO_2_ in air at a relative humidity of 80 %.

### 2.5 Phenotype of cells

Spinal cord cells were phenotyping using flow cytometry and immunohistochemistry. To perform flow cytometry studies, cells were plated into 6 wells culture plates and incubated for 48 h under the conditions described above. After 48 hours, cells were trypsinized and adherent cells combined, washed with phosphate-buffered saline (PBS: 137 mM NaCl and 2.7 mM KCl in 10 mM phosphate buffer at pH 7.4), counted and adjusted to 7×10^5^ cell/1000uLs. The cells were permeabilized by treatment with a 0.03 % (v/v) solution of saponin in PBS for 30 min at room temperature. Following incubation, cells were washed 3 times with PBS for 5 min and centrifuged at 1500 rpm. The primary antibodies used were: mouse monoclonal anti-glial fibrillary acidic protein (GFAP) (1:400; Sigma) and goat polyclonal anti-ChaT (1:100; Millipore). The secondary antibodies used were: goat anti-mouse IgG Alexa Fluor® 488 (1:1000, Invitrogen) and donkey anti-goat IgG Alexa Fluor®488 (1:1000, Invitrogen). The samples were read for FACScalibur Cytometry and analyzed by FCS Express V3.

For immunohistochemical assays, cells were plated and fixed with 2 and 4 % formaldehyde solution in phosphate-buffered saline (PBS), pH 7.4. The cells were permeabilized by treatment with 0.1 % (v/v) solution of Triton X-100, 2 % BSA in PBS. They were incubated overnight at 4 °C with the primary anti-ChaT and anti-GFAP antibody diluted to 1/100 in a 2 % solution of BSA in PBS. The secondary antibody Alexa anti-mouse IgG Alexa Fluor® 488 (1:1000, Invitrogen) and donkey anti-goat IgG Alexa Fluor®546 (Invitrogen) was diluted to 1/200 in a 2 % BSA solution in PBS and incubated for 2 h.

### 2.6. Flow Cytometry

The quantity of 5×10^4^ of cells (in 100 μL) was used per test sample and stained with PI/Annexin-FITC Apoptosis Kit solution as described previously (Matias et al., 2016, 2013). Cells were stained for 1 hour at 37 °C in the tissue culture incubator, protected from light exposure. After three washes with 1X Apoptosis Wash Buffer and added 100 μL of Wash Buffer. Samples were stained for 10 minutes at room temperature and then resuspended in 200 μL of Wash Buffer. Cell samples were ready for data acquisition on the FACScalibur.

### 2.7. Extraction of nuclei and Western Blotting

Nuclei from spinal cord cells were incubated in the presence or absence of H_2_O_2_ (100 μM) as described above, were isolated according to (Lago et al., 2017). Briefly, cells were inoculated into culture bottles containing appropriate medium (150 cm^2^ surface area of culture) and incubated for 3 and 6 hours at 37 °C in an atmosphere of 5 % CO_2_ in air at a relative humidity of 80 %. Following incubation, cells were trypsinized and adherent cells combined, washed with PBS, and centrifuged (250-300 x *g*, 10 minutes, 4 °C). The pellet resuspended in 500 μL of lysis buffer (10 mM NaCl, 3 mM MgCl_2,_ and 0.5 % Tergitol NP-40 in 10 mM Tris buffer at pH 7.5) and left on ice for 5 minutes. Cells were subsequently centrifuged (500 x *g*, 5 min, 4 °C) and the pellet resuspended in 500 μL of lysis buffer and re-centrifuged. The pellet from the second centrifugation (containing extracted nuclei) was resuspended in 150 μL RIPA buffer (150 mM NaCl, 5 mM EDTA, 1 mM dithiothreitol, 1 % Triton X-100, 0.5 % sodium deoxycholate and 0.1 % SDS in 50 mM Tris at pH 7.5) containing protease inhibitor cocktail for mammalian cells (Sigma) and centrifuged (14000 x *g*, 20 min). Supernatants and pellets were transferred to new Eppendorf tubes, and extracts were submitted to SDS-PAGE and blotted onto nitrocellulose membranes (GE Healthcare Life Sciences) with the loading of protein being confirmed by internal nuclei and cytoplasm fraction control blotting of Lamin A, GAPDH (ABCAM) and β-actin antibodies (Sigma). Membranes were blocked for 1 hour in blocking solution comprising 5 % nonfat-dried milk (Sigma) and 0.0025 % sodium azide solubilized in TBS-T (150 mM NaCl, 50 mM Tris at pH 7.5 and 0.05 % Tween-20), and washed twice with TBS-T. The primary antibodies employed were mouse monoclonal anti-GAPDH (Sigma), rabbit polyclonal anti-Lamin A (ab26300 Abcam), rabbit polyclonal anti-SOD1 (ab13498 Abcam), mouse monoclonal anti-β-actin (Sigma). The specific protein complexes formed following treatment with specific secondary antibody (anti-mouse or anti-rabbit IgG-peroxidase conjugate) were detected using SuperSignal West Pico chemiluminescent substrate.

### 2.8. Statistical analyses

All experiments were repeated at least three or five times (except where stated otherwise), and the results are expressed as the mean values ± standard deviations. Analysis of variance (ANOVA) with the Bonferroni correction was used to evaluate the differences between means, and the level of significance was set at p<0.05. The statistical analyses were performed by GraphPad software.

## 3. Results and discussion

The role of SOD1 enzyme mediating oxidative conditions in neuropathologies was investigated in spinal cord mixed culture cells. We used the spinal cord cell model for the development due to its recognized medical importance in neurobiology. This model is well known as a central nervous system development model, glial and neuronal cell interactions, ion channels, signal translation in neurodegenerative diseases, and cognition and memory formation.

The study of a collaborative tissue microenvironment *in vitro* is always challenging but also necessary to elucidate biological processes more cleanly. Here we took advantage of the spinal cord rats’ tissues explants and primary culture to explore an ALS-like microenvironment. The spinal cord is composed of neurons, microglia, and astrocytes. To perform cellular characterization of this mixed cell culture, we used the markers Choline acetyltransferase (ChaT) and glial fibrillary acidic protein (GFAP) to define motor neuron cells and astrocytes, respectively. After spinal cord dissociation, flow cytometry and immunohistochemistry techniques characterized the primary culture. In general, two populations of cells were present in the spinal cord neurons and astrocytes. The flow cytometry results in a primary population of astrocytes 12.3% with GFAP (astrocytes) and the presence of 5.6% ChaT positive cells a motor neuron marker (data not shown). The immunofluorescence staining also demonstrated a main population within the co-culture composed by astrocytes (13% of GFAP positive cells) (Figure 1C,D) concomitant to a smaller population of motor neurons (3% of ChaT positive cells, Figure 1E,F) of the total cell population. Astrocytes are essential players in both maintenances of motor neuron homeostasis and ALS disease progression, as reported before (Allen et al., 2019; Trias et al., 2018). Astrocytes play an important role in neurodegenerative disorders, as shown in human stem cells engrafted in rat spinal cord mechanically injured. Grafts also expressed GFAP by 3 months post-grafting followed by ChaT and glutamate transporters detections proving the restoration of neuron-synapse (Kumamaru et al., 2018).

**Figure 1:**
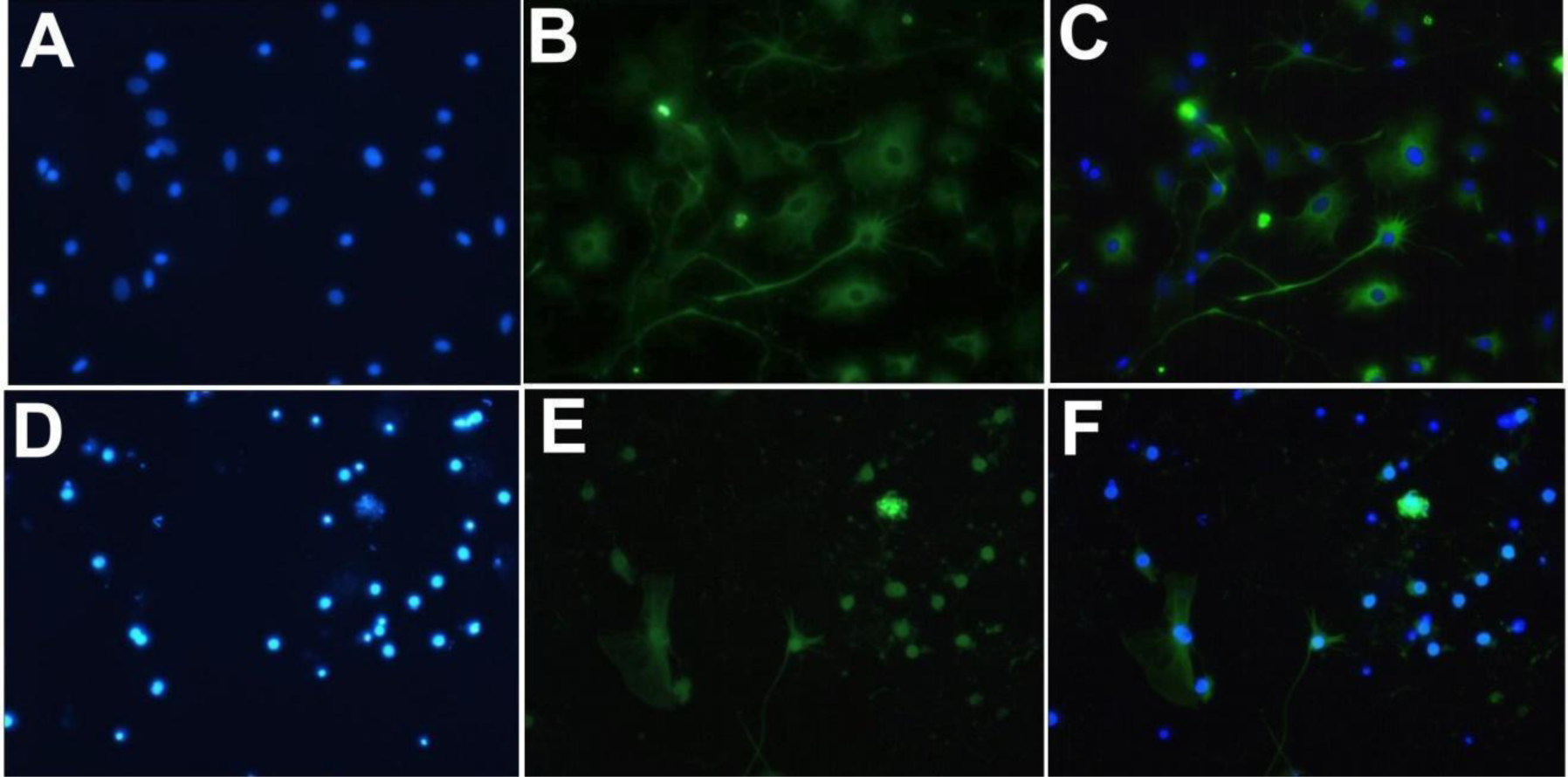
Representative micrograph showing immunohistochemistry characterization of mixed cell populations obtained by cultivation of spinal cord explants primary culture. A) Nuclei of cells labeled with DAPI, B) Cells labeled with Anti-GFAP ; C) Overlapping images showing astrocyte marker. D) DAPI, E) Cells labeled with Anti-ChaT, F) Image overlap showing the motor neuron markers.

To assess the influence of oxidative stress as neurodegenerative disease models, we treated the primary spinal cord cultures with 100 μM H_2_O_2_ during 3 and 6 hours of incubation, and flow cytometry was employed to analyze viable and dead cells. The results of the activation of cell death in spinal cord culture showed that cells treated with 100 μM H_2_O_2_ presented a decrease in the percentage of cells in early to mild apoptotic stage and an increase in dead cells when compared to control cells. Cells treated for 6 hours with 100 μM H_2_O_2_ also showed a decrease in the percentage of cells in the early apoptosis phase, but an increase in the proportion of dead cells and late apoptosis (Figure 2).

**Figure 2:**
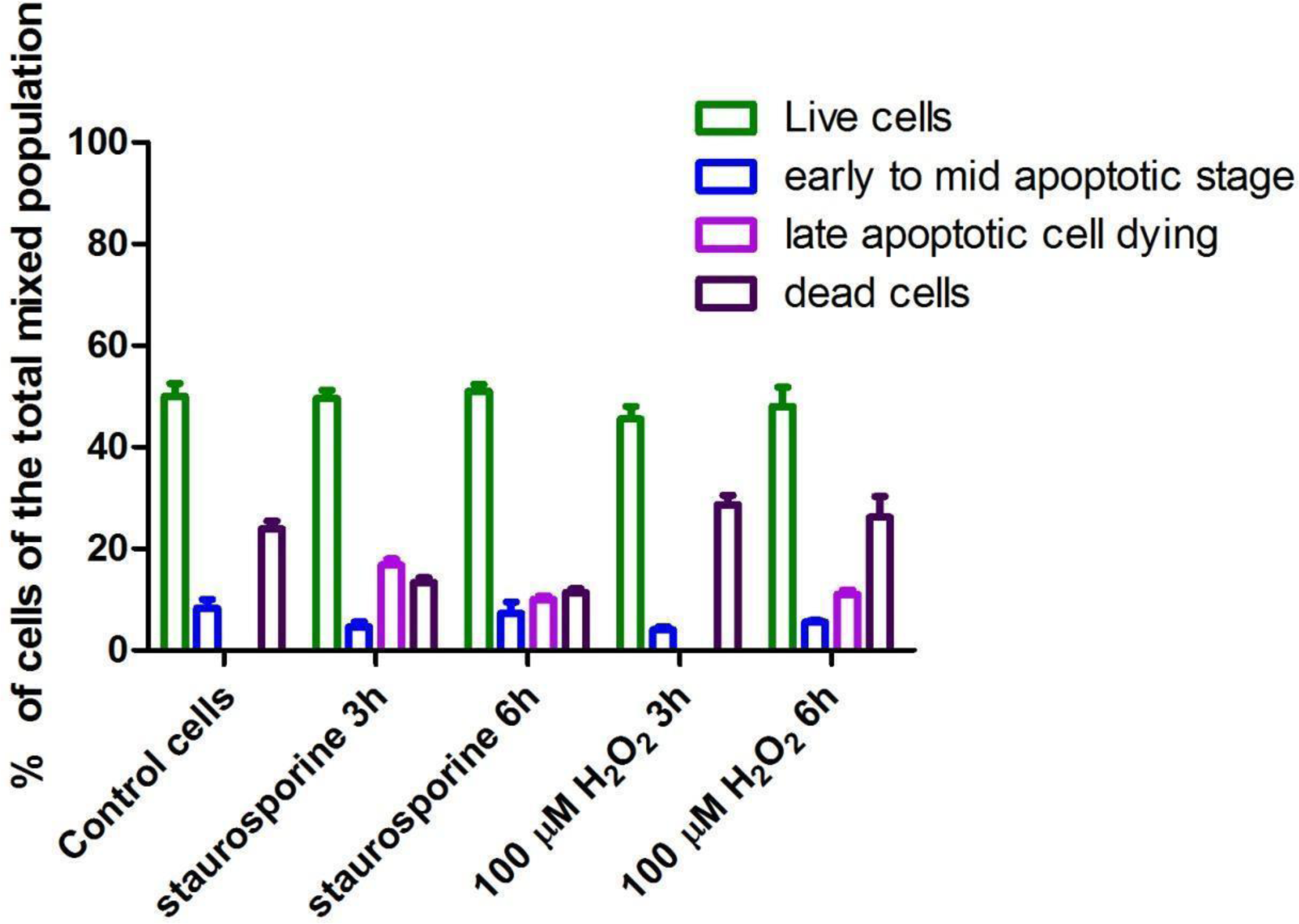
Flow Cytometry analysis of caspase cascade activation and cell death activation in primary culture cells of the spinal cord. Cells were treated with 100 μM H_2_O_2_ for 3 and 6 hours, labeled with SR-VAD-FMK and Caspase 7-AAD and compared. Data represent the mean values±standard deviations (n=3), and significant differences between untreated and treated cells were ***P ≤0.01 and **P ≤0,1.

Superoxide dismutases (SODs) are the major antioxidant defense systems against superoxide, which consist of three isoforms of SOD in mammals: the cytoplasmic Cu,ZnSOD (SOD1), the mitochondrial MnSOD (SOD2), and the extracellular Cu,ZnSOD (SOD3). Evidence suggests that in each subcellular location, SODs catalyze the conversion of superoxide to H_2_O_2_, which may participate in cell signaling (Fukai and Ushio-Fukai, 2011). Milani et al. demonstrated that the accumulation of reactive oxygen species (ROS) and increased SOD1 gene expression within SH-SY5Y cells treated with H_2_O_2_ presented pathological features of ALS (Milani et al., 2013). Other studies showed that oxidative stress regulated SOD1 transcriptionally (Dell’Orco et al., 2016). Studies indicated that SOD1, an enzyme predominantly cytoplasmic, can move to the nucleus when cells are exposed to conditions of oxidative stress, and this change of location of SOD1 showed effects against DNA damage (Tsang et al., 2014). Therefore, to explore possible cell translocation of SOD1 by oxidative stress, we treated the spinal cord mixed-cells in culture with the 100 μM H_2_O_2_ for 3 or 6 hours. We analyzed the separated nuclei and cytosol regarding the expression levels of SOD1 in each fraction by Western blotting (Figure 3), using LaminA, and GAPDH as nuclei and cytoplasmatic markers, respectively.

**Figure 3:**
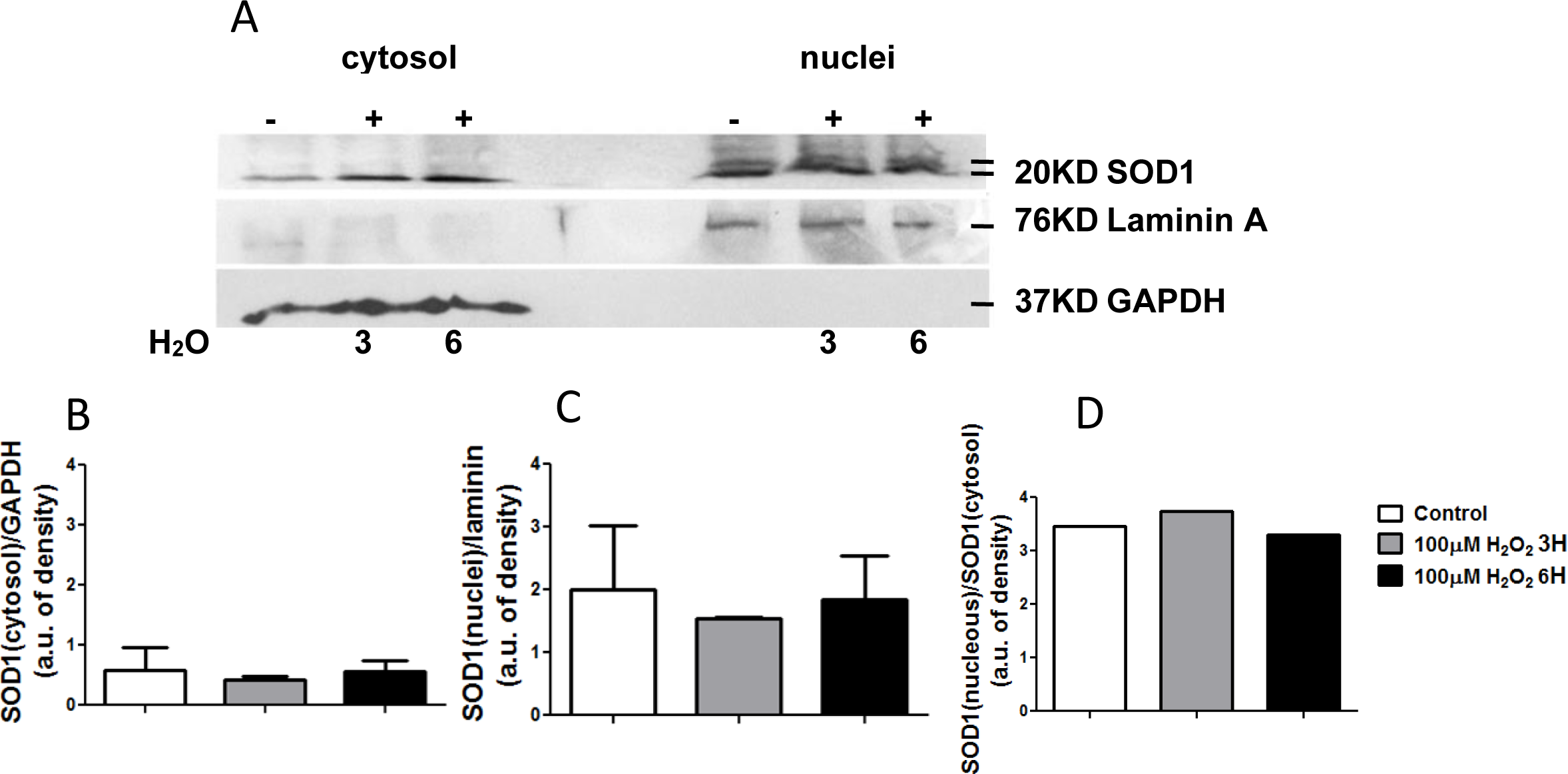
SOD1 levels in nuclei and cytosol fraction of primary culture of the spinal cord cells treated with 100 μM H_2_O_2_ and in untreated cells, after 3 and 6 hours of incubation. Lamin A was used as nuclear control; GAPDH was used as cytoplasmatic control. (A) Western blots are from one experiment and represent three identical replicates; (B) GAPDH, (C) Lamin A and (D) SOD in cytosol were used as a loading control, and densitometry of each lane (represented as bars) was calculated using the Quantity One software. The data are expressed as arbitrary units of each protein and represent the mean ± SD of *n* = 3 independent experiments, \**p* < 0.01; \*\**p* < 0.001.

The cells treated with H_2_O_2_ showed an increase in the expression level of SOD1 in their cytosolic fractions when compared to their respective controls. In the case of primary spinal cord-mixed culture (Figure 3), we can observe the translocation of SOD1 in the nucleus even in the control cells, without hydrogen peroxide oxidative treatment. This fact suggests that these cells were already submitted to oxidative stress, as we can observe in the fraction of dead cells by flow cytometry (Figure 2, control cells). Therefore, these results suggest that the translocation of SOD1 to the nucleus minimizes the effects of oxidative stress induced by hydrogen peroxide, or basal oxidative stress that we found in primary cultured cells from newborn rats. As a result, we speculate that SOD1 translocation to the nucleus in primary cells is essential to minimize the oxidative stress effects in neuronal cells or as a protection in early development.

## 4. Conclusion

Hydrogen peroxide is not an inducer of SOD1 activity. Still, the translocation of SOD1 - a cytoplasmic enzyme - to the cell nucleus under oxidative stress caused by H_2_O_2_ in model cells in culture has been observed in the literature. Based on our results, when submitting mixed spinal cord culture cells from newborn rats containing motor neurons and astrocytes to the oxidative stress treatment, we observed the translocation of SOD1 to the cell nucleus even without the presence of oxidative stress. In this case, the conclusion that only cells submitted to the presence of H_2_O_2_ exhibit the SOD1 nuclear translocation cannot be reached. We observed the occurrence or presence of a high basal oxidative stress in spinal cord cells of neonatal rats or another mechanism that does not involve oxidative stress in the translocation of SOD1 to the cell nucleus, requiring further investigation into this phenomenon.

## Acknowledgments

This work was supported by the Brazilian agencies São Paulo Research Foundation (FAPESP, Grant 2018/14152-0 and 2020/14175-0), CAPES (Financial code 001), UFABC Multiuser Equipment Facilities, and the National Council for Scientific and Technological Development (CNPq). NVS and ACM thanks FAPESP for fellowships during their PhDs.

## Author Contributions

GC designed the study. CMLM, NVS and GC performed the experiments. GC and CMLM analyzed the data and wrote the paper.

## Notes

### Competing Interest Statement

The authors have declared no competing interest.

